# Megaloneurite, a giant neurite of vasoactive intestinal peptide and nitric oxide synthase in the aged dog and identification by human sacral spinal cord

**DOI:** 10.1101/726893

**Authors:** Yinhua Li, Wei Hou, Yunge Jia, Chenxu Rao, Zichun Wei, Ximeng Xu, Hang Li, Fuhong Li, Xinghang Wang, Tianyi Zhang, Jingjing Sun, Huibing Tan

**Affiliations:** Department of Anatomy, Jinzhou Medical University, Jinzhou, Liaoning 121001, China; Key Laboratory of Neurodegenerative Diseases of Liaoning Province, Jinzhou Medical University, Jinzhou, Liaoning, 121001, China

**Keywords:** Megaloneurite, Giant neurite, VIP, Aging, Dog, Human, Sacral spinal cord

## Abstract

Megaloneurite of NADPH diaphorase (NADPH-d) positivity is a new kind of aging-related neurodegeneration and also co-localized with vasoactive intestinal peptide (VIP) in the sacral spinal cord of aged dog and monkey. However, no immunocytochemistry of VIP was exclusively tested in the aged dog and no evidence has been reported in the aged human spinal cord. Aged dogs were used to examine the distribution of VIP immunopositivity in the sacral spinal cord. Immunocytochemistry of VIP and alpha-synuclein were also examined in the aged human spinal cord. The VIP immunopositivity in aged dog was reconfirmed our previous finding illustrated by immunofluorescent study. Megalogneurite was also identified by nitric oxide synthase (NOS) immunoreaction in aged dog. The VIP positive megaloneurites both in age dog and human were detected in dorsal root entry zoon, Lissauer’s tract, dorsal commissural nucleus and anterior commissural gray as well as in the lateral funiculus of the sacral spinal cord exclusive of other segments of spinal cord. Alpha-synuclein positivity was present mini-aggregation and Lewy body in the sacral spinal cord of aged human, that also occurred in the lumber, thoracic and cervical spinal cord. It was firstly tested that VIP megaloneurites occurred in the aged human sacral spinal cord, especially in the white matter. Megaloneurites identified by NADPH-d-VIP-NOS immunoreaction could implicate for the dysfunction of pelvic organs in the aged human being.

---

Vasoactive intestinal peptide (VIP) acts on neuronal cells to support physiological condition and protect neurons survival in pathological condition[1-3]. Different from other neuropeptides, the distribution of VIP is restricted to the sacral cord[4]. Or it is majorly distributed in the sacral spinal cord and a few of VIP positivity is detected in the other segment of the spinal cord in cat[5].VIP neurons are small-size neurons and central original source for peripheral innervation[6]. VIP-ergic afferent neurons projected to the pelvic organs are bigger proportion that the other segments, while the lower lumbosacral segments of the neuropeptide innervation was also more priority to the upper lumber segment[7]. VIP fibers are majorly found in the Lissauer’s tract (LT), superficial lamina, the intermediate gray, and the central gray dorsal and around central canal[8]. Chung reported that many VIP fibers are detected as thick fascicles in LT and in the intermediate gray[8]. But detailed distribution pattern and associated properties with specialized terminology is still lack. We thought that colocalized with NADPH-d positive megaloneurites[9, 10]. The verification of megaloneurites by VIP examination should be considered in sacral spinal cord of aged human, because distribution of VIP is significantly higher in dorsal than in ventral gray matter in human[11]. Compared with several other neuropeptides, the concentration of VIP shows much higher in the lumbosacral spinal cord in human[12, 13]. Anand suggested: “some of VIP-like immunoactivity is the primary afferent fiber” in his study. Pelvic organs are functionally innervated by VIP-ergic neurons[14-19]. For example, VIP-ergic neurons and fibers act potentially on erectile function in both of dog[20-22] and human[16, 23]. VIP is related to age in patients of pelvic organ prolapse[24]. Compared with VIP, several preproVIP-derived peptides did not cause a significant inhibition of smooth muscle activity of human female genital tract [25]. VIP also plays important role in somatic pain sensation[26]. Somatic nerve injury causes upregulation of VIP in the sacral spinal cord, while pelvic nerve injury shows downregulation[27].

NADPH diaphorase (NADPH-d) positive neuron is considered relating to nitric oxide synthase (NOS)[28]. NOS positive neurons distribute in the spinal cord[29-34] and pelvic organs[35]. In our previous study, we find aging-related NADPH-d megaloneurite in the sacral spinal cord of aged dog[9]. The similar megaloneurites are also evident in the aged monkeys[10]. The megaloneurite is colocalized with VIP[9, 10]. For human conditions and diseases, the dog recently is considered as a natural model and experimental observation of the canine aging phenotype could be concluded relating that of the human[36]. The companion dog may be used as an experimental model to study of human morbidity and mortality”[37]. However, study of VIP and NOS positivity in the aged dog is still needed to confirm in the aged dog. In the present study, VIP and NOS immunoreactivity in aged dog is examined to test megaloneurite in aged dog. Next, we also test the distribution of VIP positivity for megalonueirte in aged human sacral spinal cord.

## Materials and methods

Domestic dogs (Canis lupus familiaris) of young adult (n=6, 1∼ 5-year-old), aged (n=6, ≥10-year-old) dogs (6– 15 kg) and 8 human spinal cords (79∼108-year-old) of both sexes from donated cadavers for medical education program were used in the present experiments. The animals were anesthetized with sodium pentobarbital (30-50 mg/kg i.v.) and perfused transcardially with saline followed by 4% paraformaldehyde in a 0.1M phosphate buffer (PB, pH 7.4). After perfusion fixation, the spinal cords were rapidly obtained and followed in 30% sucrose for 48 hrs. The spinal cords from the cervical to coccygeal segments were sectioned by 40 μm thickness on a cryostat. Human spinal cords fixed with 10% formalin were followed in 30% sucrose for 48 hrs and cut transversely by 40 μm thickness on the same cryostat.

### NADPH-d histochemistry

Staining was performed using free floating sections (Tan et al., 2006). Most of the spinal cord sections from the young and old dogs were stained and examined by NADPH-d histochemistry, with incubation in 0.1 M Tris-HCl (pH 8.0), 0.3% Triton X100 containing 1.0 mM reduced-NADPH (Sigma, St. Louis, MO, USA) and 0.2 mM nitro blue tetrazolium (NBT, Sigma), at 37°C for 2 to 3 h. Sections were monitored every 30 min to avoid overstaining. The reaction was stopped by washing the sections with the phosphate buffered saline (PBS, 0.05M).

### Immunocytochemistry

Some dog sections were mainly processed VIP, nitric oxide synthase (NOS) immunocytochemistry, respectively. Human sections were processed VIP and α-synuclein immunocytochemistry. The sections were collected in PBS in 24-well plates and processed for free-floating immunocytochemistry using primary polyclonal antibodies of VIP (rabbit, 1:1000 Sigma, USA) NOS (Cayman Chemical,1:1000, USA), neuropeptide Y (NPY, rabbit; 1:5000, Sigma, USA), calcitonin gene-related peptide (CGRP, mouse; 1:100, Sigma, USA) or α-synuclein (rabbit; 1:100, Sigma, USA). For inactivation of the endogenous peroxidase, free-floating sections were rinsed with 0.3% hydrogen peroxide in methanol for 10 minutes. Sections was then washed with 0.1 M PBS (pH 7.2) for 30 minutes, and immunostained by means of the Strept(avidin)-Biotin Complex (ABC) method. Briefly, 0.1% Triton X-100 was added to the immunoreagents to increase antibody penetration. The sections were rinsed in antibody solution (0.1% Triton X-100, 3% normal goat serum / 3% bovine serum album in PBS) for 30 minutes and incubated in primary antisera overnight at room temperature (16-24 hours). The dilutions of antibodies were 1:1,000 for VIP, and 1:1,000 for NOS. The primary antibody was replaced by antibody solution (30 minutes), followed by antibody solution (30 minutes) and then incubated in a biotinylated secondary antibody (1:800) at room temperature (1 hour). The tissue was rinsed in antibody solution for 30 minutes each and incubated in a streptavidin–enzyme complex (1:800 in PBS) at room temperature (1 hour). After rinsing in PB for 30 minutes, sections were incubated in 3,3-diaminobenzidine hydrochloride (DAB, Sigma; 0.05% in 0.1 M PB, pH 7.2) and 0.01% hydrogen peroxide (5-10 minutes). Antibody-based avidin-biotin reagents (VECTASTAIN® ABC system) were purchased from Vector Laboratories. In each immunocytochemical testing, a few of sections were incubated without primary antibody, as a negative control. Finally, the sections were mounted on slides and coverslipped. For controls of immunocytochemistry, the primary antibodies were replaced with the same amount of normal serum from the same species while doing the same specific labeling with the normal procedure of immunocytochemistry. No specified staining was detected in the immunostaining control procedure.

## Statistics and Figure edition

Statistical analyses were performed using GraphPad Prism 5.0 (GraphPad Software, La Jolla, CA). Differences between human and aged dogs of VIP positive segments of megaloneurite in lateral funiculus of the sacral spinal cord were analyzed by using student’s *t*-tests. Diameter of VIP positive fiber and megaloneurite was analyzed by the same statistics test. All data are indicated as the mean ± SEM and P < 0.05 were regarded as statistically significant.

## Results

NADPH-d positivity revealed both NADPH-d positive neuronal soma and fiber in the spinal cord. The results were consistent with the previous study[38]. NADPH-d positive megaloneurites were detected in the sacral spinal cord of aged dog (Figure 1). No megaloneurite was detected in the other segments of spinal cord. The results were also consistent with our previous study[9]. Our next examination was to define and verify if any other chemical or neuropeptide neuronal similarity was revealed in the sacral spina cord of aged dog. VIP positive megaloneurites occurred in the sacral spinal cord of aged dog (Figure 2). The megaloneurites were distributed in the dorsal horn, dorsal root entry zone (DREZ), LT, dorsal gray commissure (DGC), lateral collateral pathway (LCP), medial collateral pathway (MCP) and area around central canal. Different to examination of NADPH-d staining, we did not find VIP positive neuronal soma in both young and aged dog. VIP VIP megaloneurites were dilated-diameter fibers of no clear varicosity (Figure 3). In the area of the central canal, some megaloneurites penetrated between ependymal cells into lumen of the central canal. The NOS immunoreaction revealed megaloneurites in the aged dog (Figure 4). Beside existence of NOS megaloneurites in gray matter, there were NOS megaloneurites in the dorsolateral funiculus (Figure 4D). Next, neuropeptide Y immunoreactivity was examined in both young and aged dog. No megaloneurite was detected by the neuropeptide Y immunoreaction in the sacral spinal cord of aged dog (Figure 5), which showed that most of immunopositivity was neuronal fibers. No megaloneurite was detected by the CGRP immunoreaction in the sacral spinal cord of aged dog (Figure 6), which similarly showed that most of immunopositivity was neuronal fibers. Finally, we examined the aged human spinal cord. VIP positive megaloneurites were detected in the sacral spinal cord of aged human (Figure 2). The VIP positive megaloneurites were distributed in the dorsal horn, DREZ, LT, DGC, LCP, MCP and area around central canal (Figure 7). The segment-like VIP megaloneurites orientated along with sensory afferent pathway in which located in DREZ, LT, DGC, LCP and MCP. Different from that of aged dog, the VIP megaloneurites also distributed to anterior commissure ventral to the central canal. The VIP megaloneurite indicated by arrowhead in in figure 7D run toward to anterior commissure. Beside distribution in the grey matter, short segments of VIP megaloneurites were detected in the lateral funiculus (Figure 8). Significant different (p<0.001) was showed between the diameter between VIP thin fiber and megaloneurite. No VIP megaloneurite was detected in the other segmental spinal cord in aged human. Different to examination of NADPH-d staining, no VIP positive neuronal soma was detected in sacral spinal cord of aged human in our examination. Different to aged dog, the number of VIP megaloneurite segments in the lateral funiculus was slightly bigger than that of aged dog and distributed area of VIP megaloneurite segments in the region was also larger than that of aged dog. The percentage area of distribution of VIP megaloneurite segments in lateral funiculus in aged human was significantly larger than that of aged dog(p<0.01). Note to compare Figure 4D to Figure 8A and Figure 8G. Summerly, beside the gray matter, megaloneurites of NADPH-d, NOS and VIP exclusively occurred the lateral funiculus of aged spinal cord.

**Figure 1.**
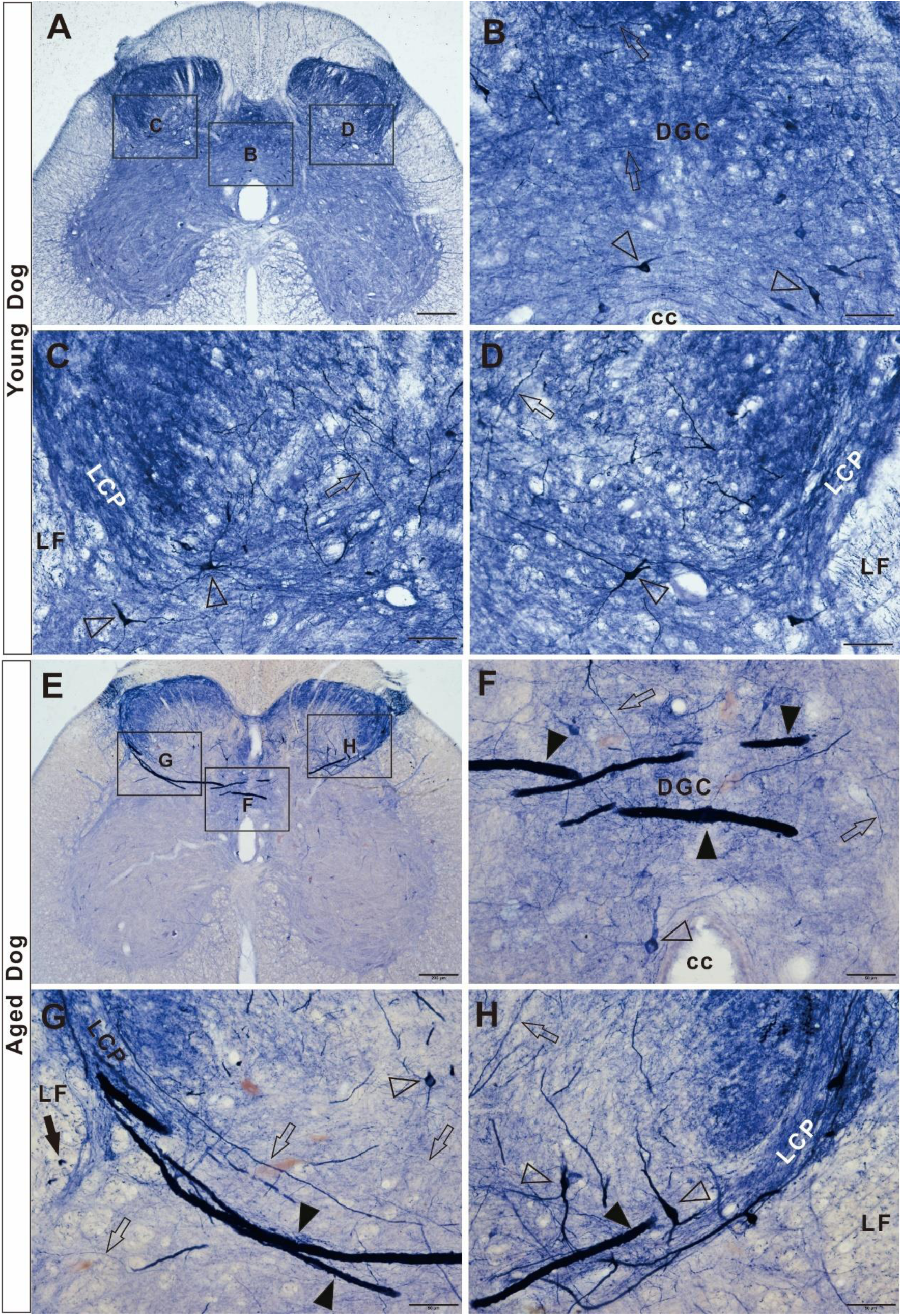
Megaloneurites detected by NADPH-d histology in the sacral spinal cord of aged dog. Arrowhead indicates megaloneurites. Open arrowhead indicates neuronal soma. Open arrow indicates regular NADPH-d positive fiber. Arrow indicates transverse of megaloneurite in lateral fusciculus (LF). CC: central canal. LCP: Lateral collateral pathway. DGC: dorsal gray commissure A-D: NADPH-d positivity in young dog. E-H: NADPH-d positivity in young dog. Megaloneurites detected in higher magnification of F-H. Bar (A and E)=200μm. The other scale bar=50μm.

**Figure 2.**
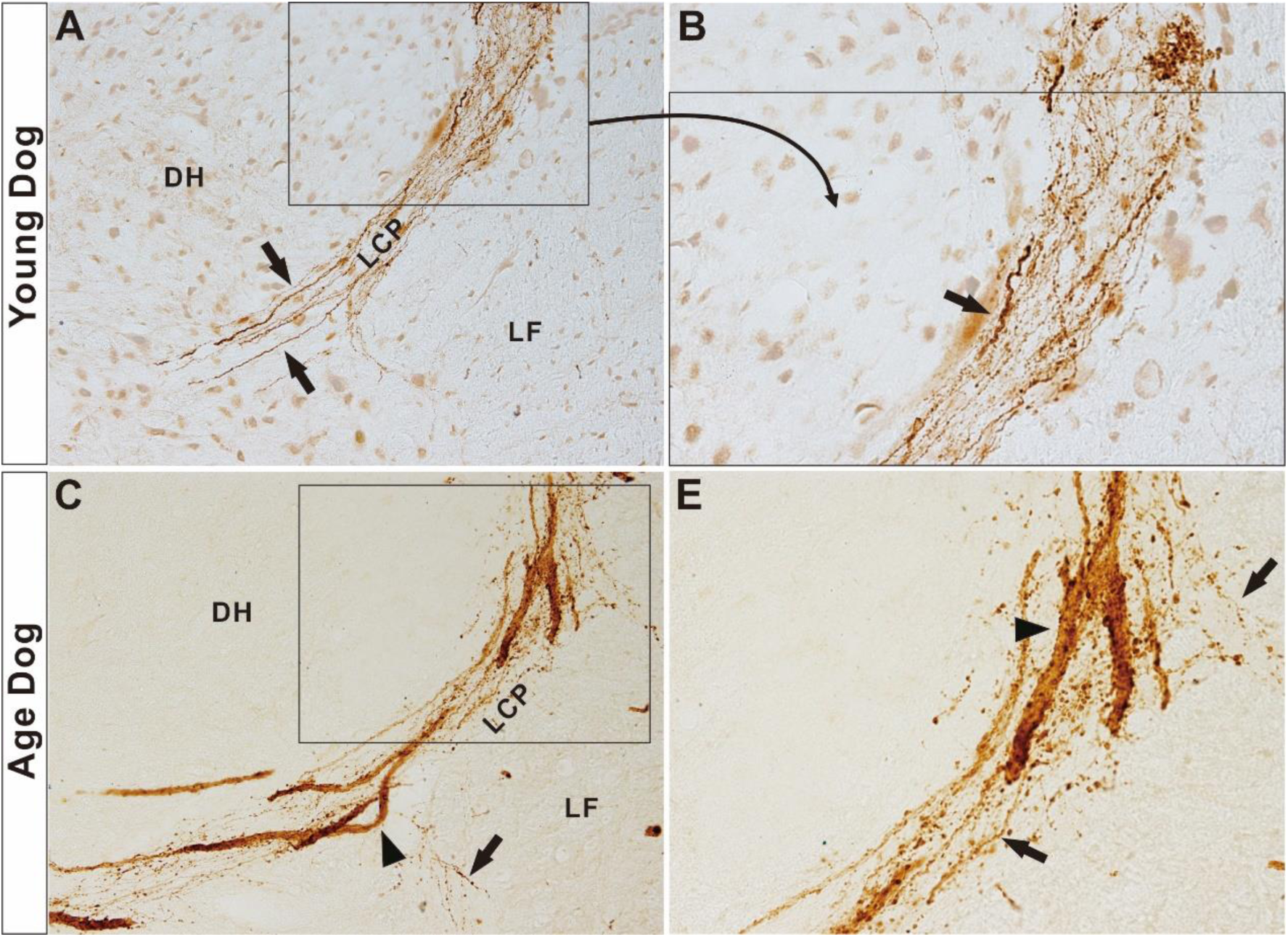
VIP positive megaloneurites in the sacral spinal cord of aged dog. DH:dorsal horn. LF: lateral fasciculus. LCP: lateral collateral pathway. A: Arrows indicate example of regular thin fibers in sacral spinal cord of young dog. B: Higher magnification of A. C: VIP positive megalonuerite (arrowhead) occurred in the LCP of sacral spinal cord in aged dog. D: Higher magnification of C showed megalonuerite (arrowhead).

**Figure 3.**
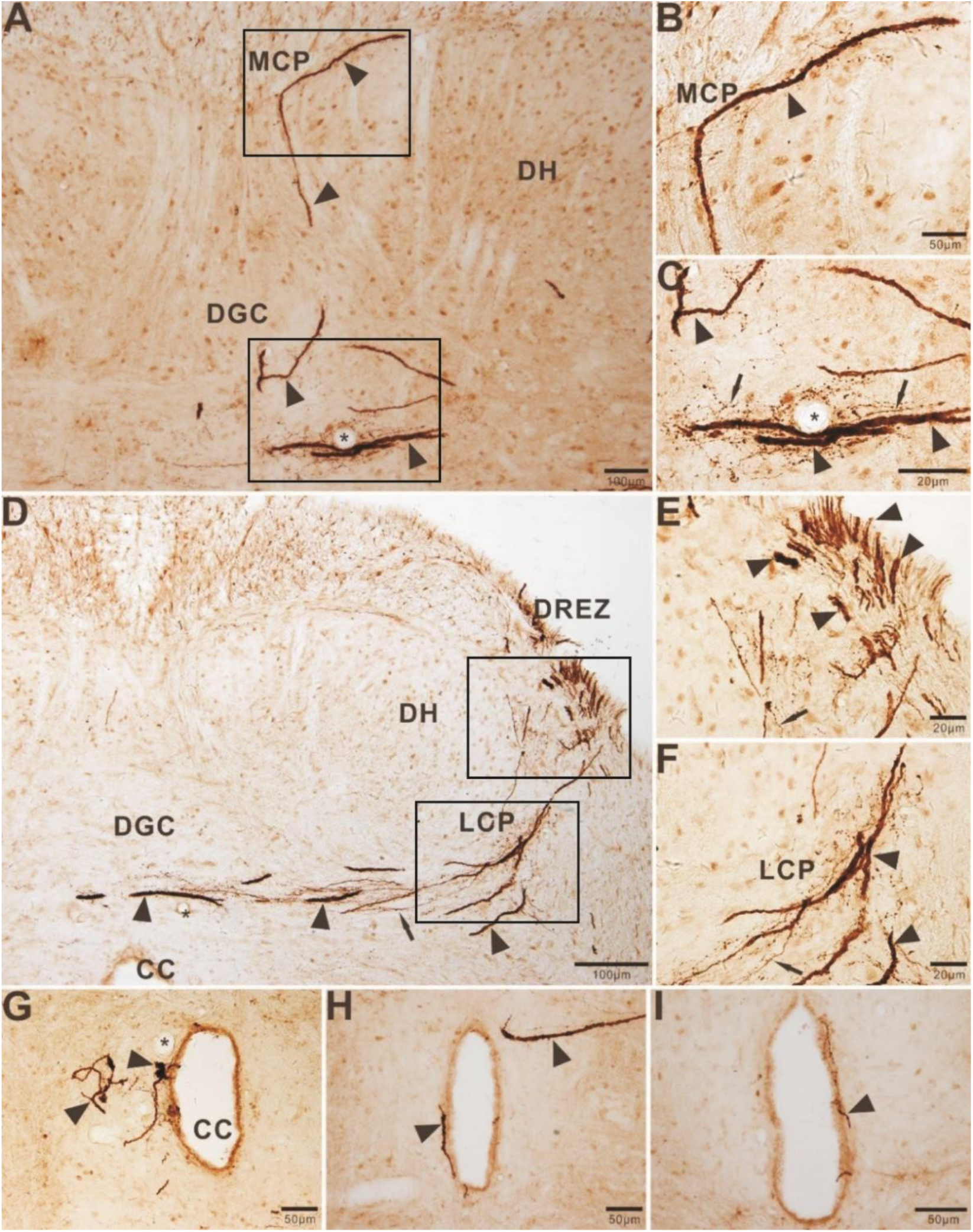
VIP immunoactivity in the sacral spinal cord of the aged dog. A-C:VIP immunoactivity showed in the dorsal commisureal nucleus, DH:dorsal horn;MCP:medial collateral pathway; DGC:dorsal gray commisture; arrowhead: megaloneurite;arrow:normal neurite;*:blood vessel. B: higher magnification from A labeled area MCP. C: higher magnification from D labeled area DCC. D-F:dorsal part of section with DREZ, LCP and DGC in the sacral spinal cord in aged dog; LCP:lateral collateral pathway;CC:central canal;DREZ:dorsal root entry zone. E: higher magnification from D labeled area DREZ. F: higher magnification from D labeled area LCP. G-I: megaloneurite distributed around area of central canal. VIP megaloneurite occurred subependymal cells(H) passed through ependymal cell reaching the lumen of CC(G&I). Bar (A&D)=100μm, B&G-I=50μm, C, E, F=20μm.

**Figure 4.**
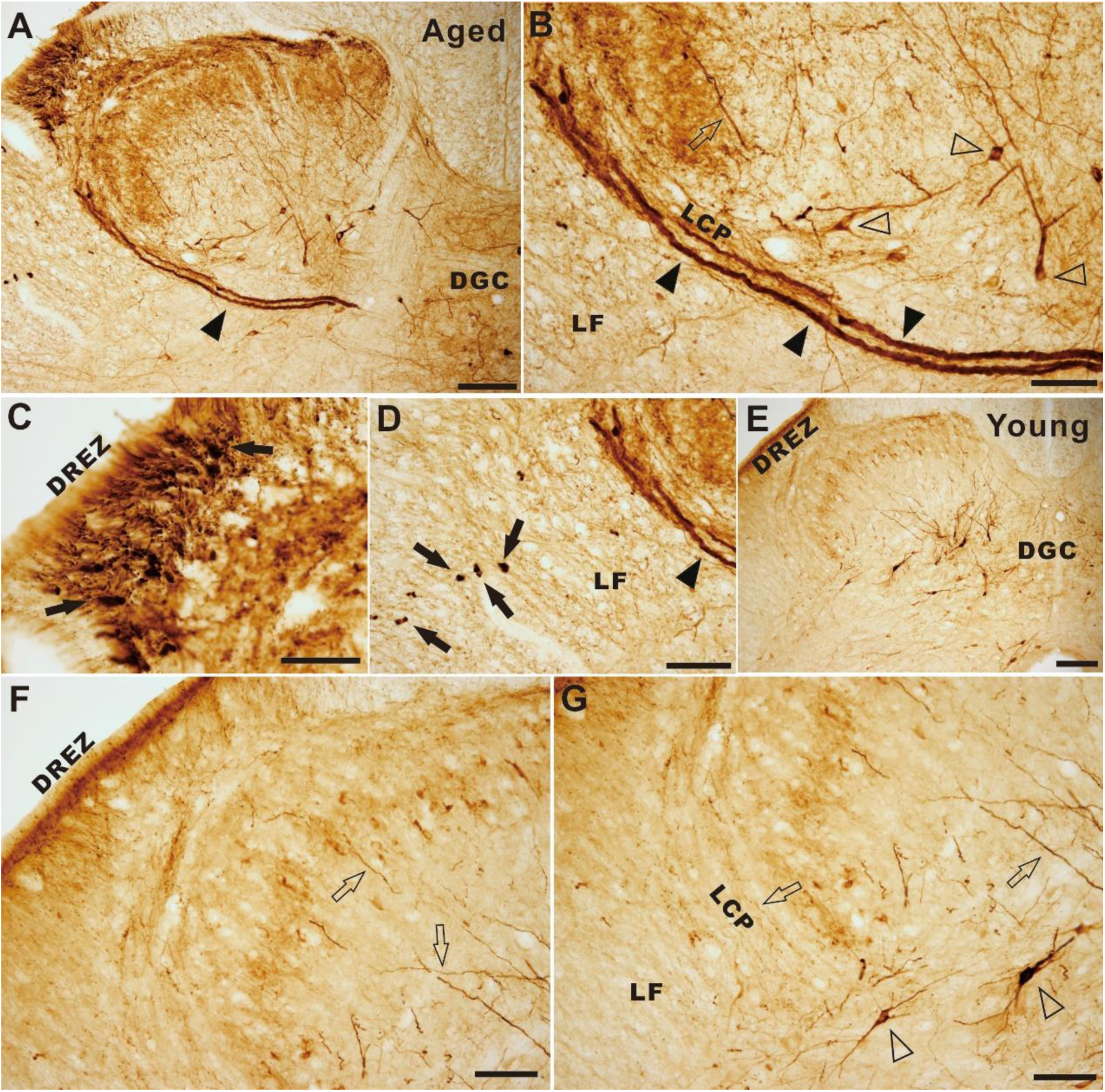
NOS positivity in the sacral spinal cord of aged and young dog. Arrowhead indicates megaloneurites. Open arrowhead indicates neuronal soma. Open arrow indicates regular NOS positive fiber. Arrow indicates transverse of megaloneurites in dorsal lateral fusciculus (LF). A: Megaloneurites (Black arrowhead) of NOS immunoreactivity in dorsal horn, lateral collateral pathway (LCP) and dorsal gray commissure(DGC)in the aged dog. B: Higher magnification image from A for LCP. C: Higher magnification image from A for DREZ. D: Higher magnification image from A for LF. E: NOS immunoreactivity in young dog. F: Higher magnification image from E for DREZ. G: Higher magnification image from E for LF and area LCP. Bar=100μm(A and E). Bar = 50μm(B, C, D, F And G).

**Figure 5.**
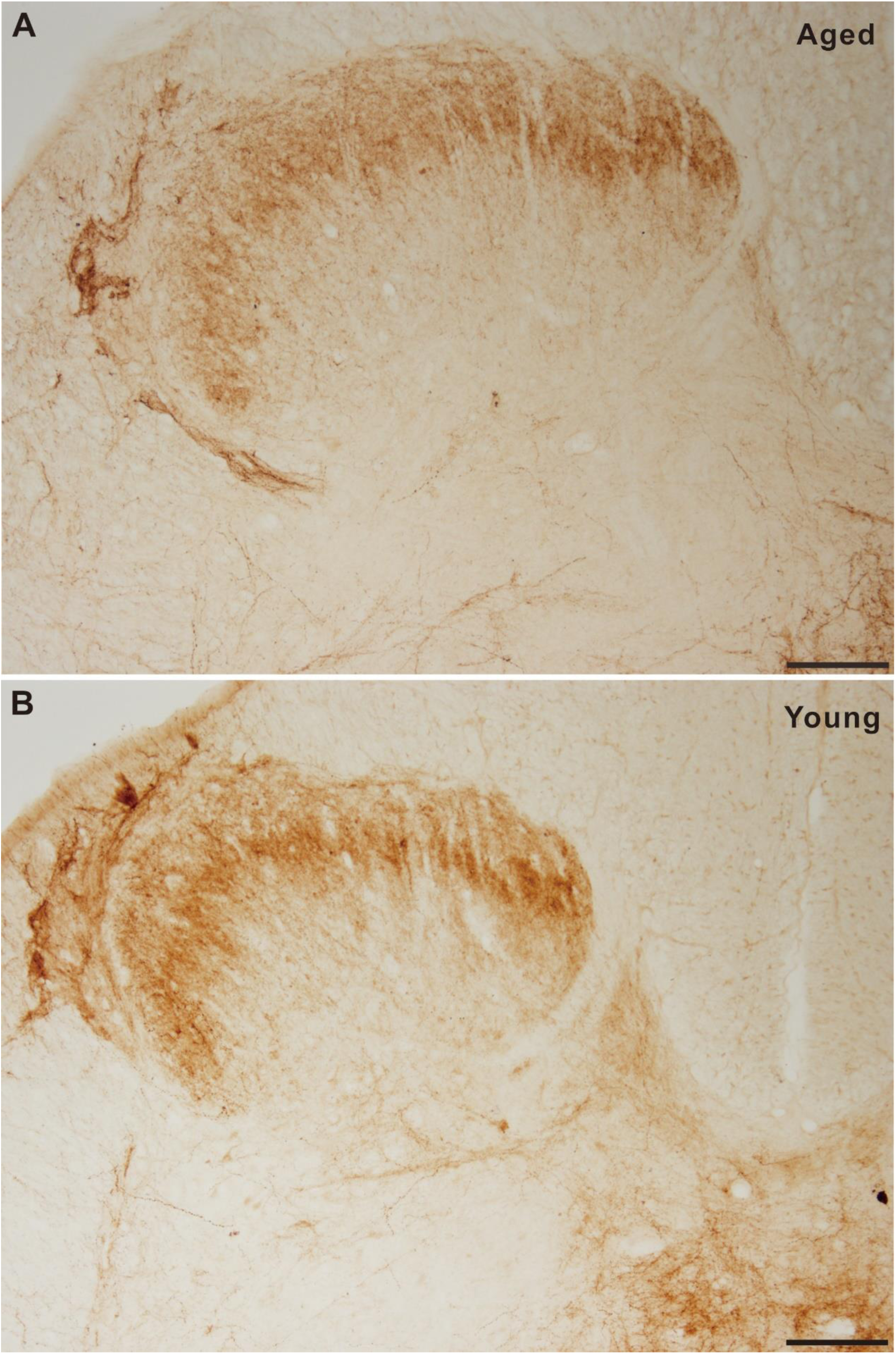
Neuropeptide Y immunocytochemistry in sacral spinal cord of aged (A) and young(B)dog. Bar =100 μm.

**Figure 6.**
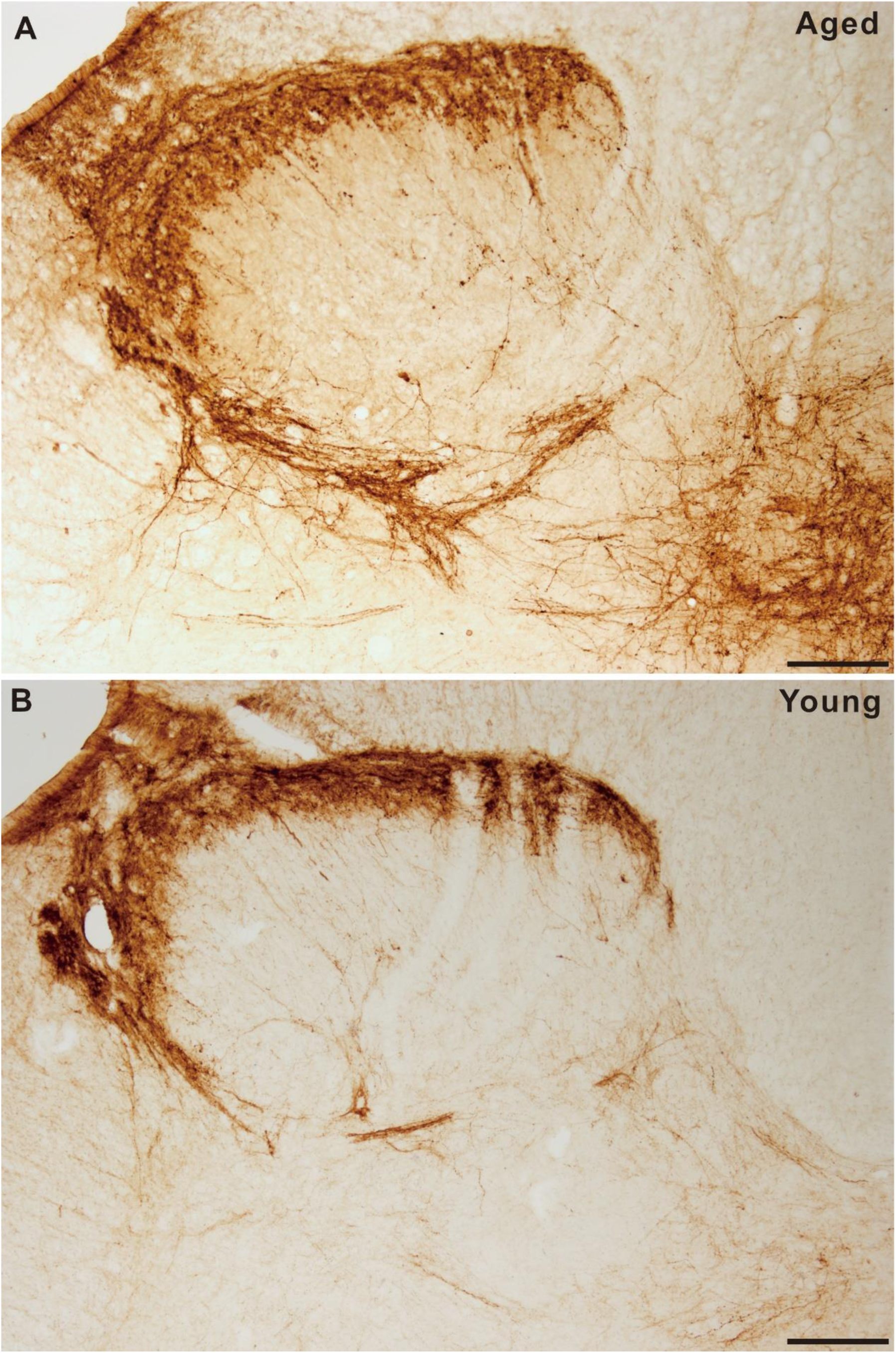
CGRP immunoreaction in the sacral spinal cord of aged (A) and young(B) dog. Bar=100 μm.

**Figure 7.**
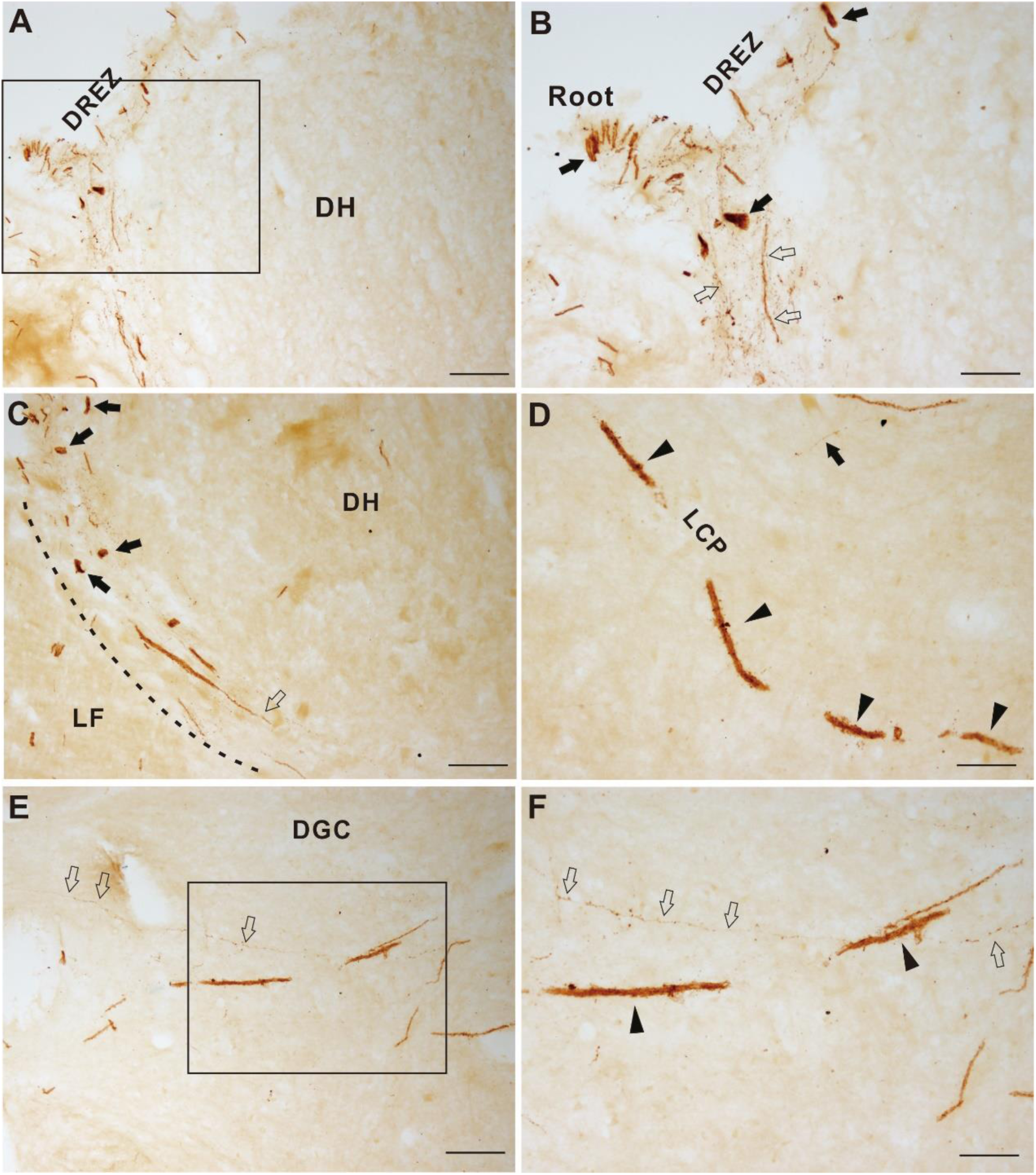
VIP megaloneurite occurred in the sacral spinal cord of aged human. Thin fibers indicated by open arrow. Arrow indicated segment of megaloneurite. Arrowhead indicated megaloneurite. A: Segmental megaloneurites detected in DREZ. B: Higher magnification of A to show segmental megaloneurites(arrow). C: Megaloneurites detected in lateral collateral pathway (LCP) and showed in higher magnification figure C (indicated by arrowhead). E: Megaloneurites detected in the dorsal gray commissure (DGC) and aslo showed in higher magnification figure C. Bar=50μm.

**Figure 8.**
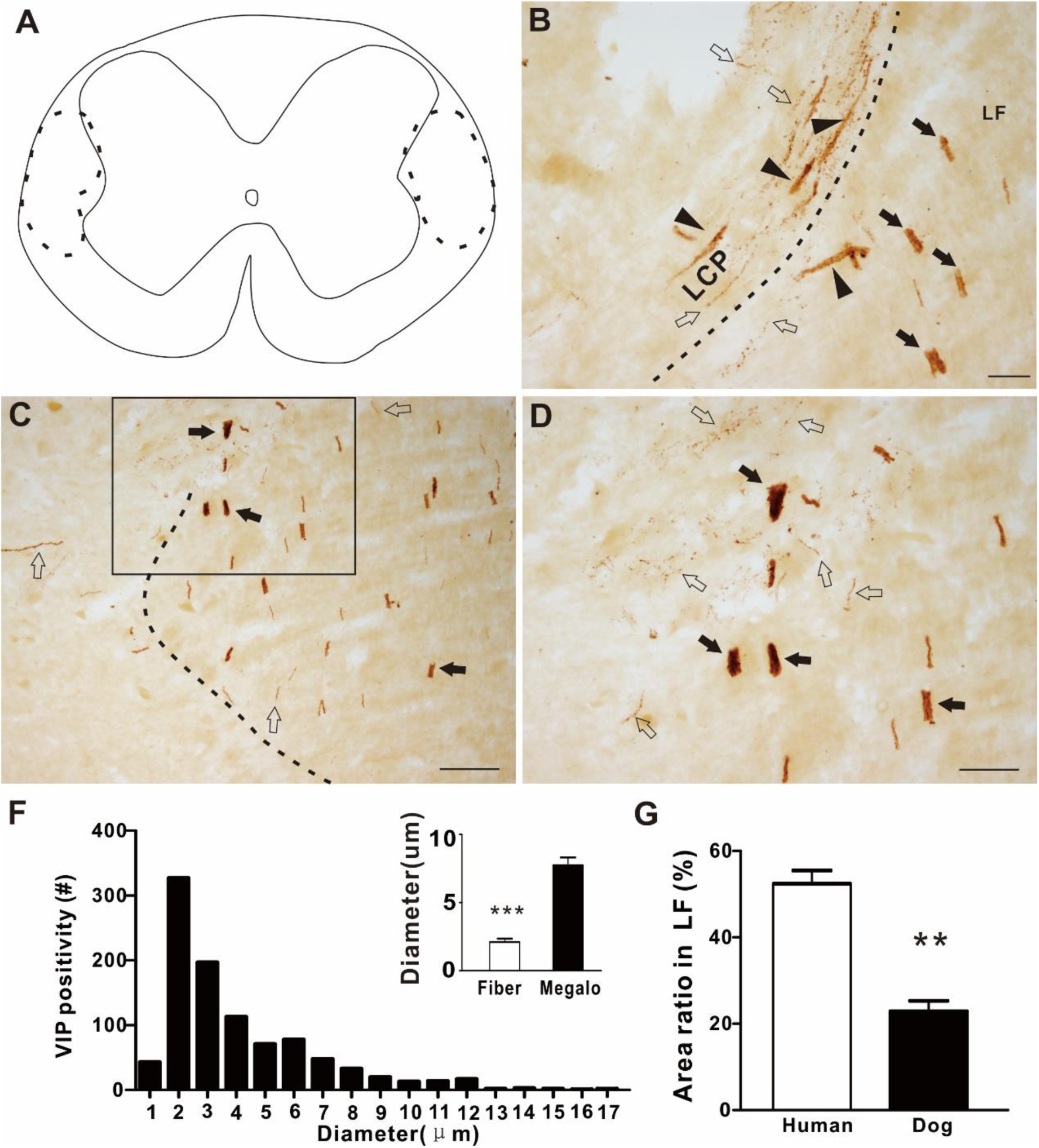
Enlarged VIP puncta detected in the lateral funiculus (lateral of LCP). A: Dashed circles indicated location of Enlarged VIP puncta in schematic illustration. Dashed lines in B and C were boundary of grey matter and white matter. Arrows indicated segmental VIP megaloneurite or enlarged VIP puncta (transverse megaloneurite). Open arrow indicated thin regular VIP fiber. D: Higher magnification of the rectangle in C. F: Histogram of diameter of 1000 VIP positive fibers and megaloneurites. Insert bar plot indicated comparison the diameter the between VIP thin fibers and megaloneurites. G: Percentage area of segments of megaloneurite in the lateral funiculus (LF) of spinal cord in aged human compared with that of aged dog. Bar =50 μm (B and D), Bar= 100μm for C.

To a certain extent, to test neurodegenerative alterations and aging deterioration relevant to sacral spinal cord[39, 40], we examined α-synuclein expression in sacral spinal cord of aged human. Mini-aggregated α-synuclein and Lewy body were detected in the LCP, DGC, anterior horn, intermediolateral nucleus and Onuf’s nucleus(Figure 9), that also occurred in the lumber, thoracic and cervical spinal cord. But no megaloneurite-like structure was detected in the sacral spinal cord of the aged human. The results of the α-synuclein immunoreactivity were similar to the previous reports[39, 41].

**Figure 9.**
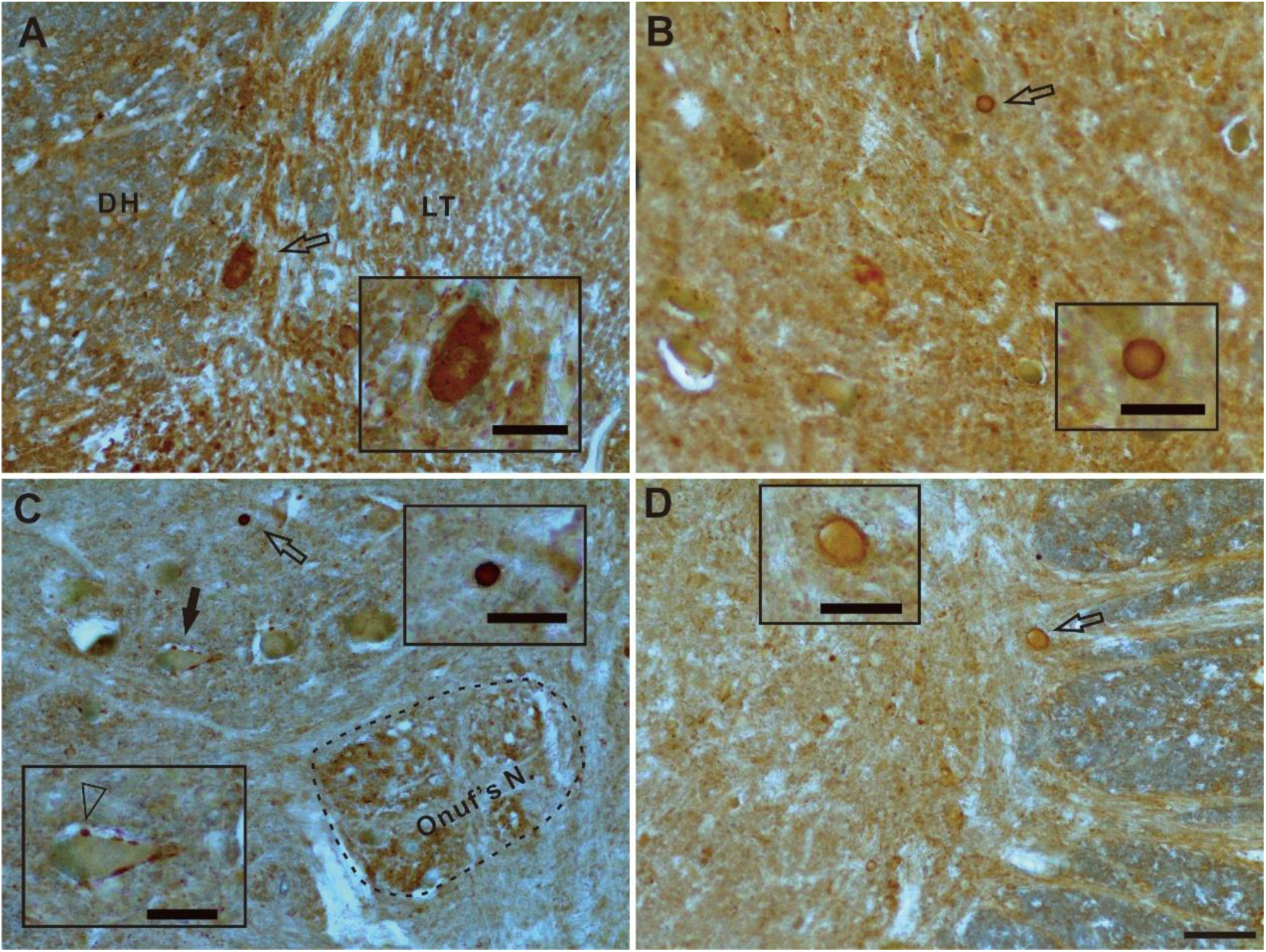
Lewy body detected by α-synuclein immunoreactivity in the sacral spinal cord of aged human. DH: dorsal horn. LT: Lissauer’s tract. Open arrow indicated Lewy body showed in insert. A:Lewy body detected in the dorsal horn. B-D: Lewy body in anterior horn. C: Arrow indicated a motor neuron surrounding mini-aggregation of α-synuclein pointed by open arrowhead. Onuf’s nucleus indicated by dash line. Bar=100μm. Bar in insert =50μm.

## Discussion

In the present study, distribution of VIP and NOS immunoreactivity was investigated by immunocytochemistry in the dog spinal cord. We reconfirmed that NADPH-d megaloneurite. Consistent with our previous investigations[9, 10], we keep using the term megaloneurites. Two types of VIP positive fibers were defined by the diameter of the neurites: thinner VIP-neurite and thicker VIP-neurite or two kinds of VIP neurites: thin varicose VIP fiber and VIP megaloneurites. NOS megaloneurites were also found in the aged sacral spinal cord. Further, VIP megaloneurites also confirmed in the sacral spinal cord in aged human. Megaloneurites distributed in the DREZ, LT, LCP, MCP, DGC and area around central canal as well as in the dorsal lateral fasciculus in the white matter in the sacral spinal cord of both aged dog and human. LCP and MCP in the sacral spinal dorsal horn are important for the relay of pelvic visceral afferents[42]. DGC is a region for integration of pelvic visceral regulation[43]. Megaloneurites occurred in the area suggested that regulation of pelvic visceral regulation could be changed for giant neurite.

It is the first time that the evidence of VIP megaloenurite was visualized in the white matter of lateral funiculus in the sacral spinal cord of aged human. VIP megaloneurites provided more detailed experimental results. The number and distribution region of VIP megaloenurites in the lateral funiculus were large than those of aged dog. We thought that NOS positive megaloneurite was important to verity the existent megaloneurite in aged sacral spinal cord. VIP is important neuropeptide to regulate pelvic organs[13, 44-46]. However, different to NO-ergic neuronal positivity, no cell body was detected in our experiments. This results is consistent with previous study[13]. Actually, we also examined neuropeptide Y and CGRP, both of the neuropeptides showed no neuronal cell body was detected in the spinal cord. Neuropeptide Y and CGRP are important neuropeptide and similar distribution of both neuropeptides[11, 12, 27, 47-49], which are relevant in pelvic afferent pathways[46]. We also tested NeuN and GFAP immunoreactivity (data not showed here). The correlation of megaloneurite will be analysis in future. NOS immunoreactivity suggested occurrence of NADPH-d positive megaloneurite. VIP immunoreaction exclusively labeled the megaloneurites in the aged dog and human in our experiments.

For terminology and concepts of megaloneurite, prefix came from following terms: megacolon[50], megalo-Ureter[51] and congenital megalo-urethra[52] as well as megalo-type α-1,6-glucosaccharides[53]. We coined “megaloneurite”, which means a giant neurite. We also noted the “giant fiber” was a specialized term for some species, for example earthworms[54], drosophila[55]. However, we misused a term “meganeurite” to define our new finding in the sacral spinal cord of aged dog, because we were thinking about “megamitochondria” and “megacolon” at very beginning without more efficient searching in internet. Actually, meganeurite is a special for neurodegeneration in lysosomal dysfunction[56] or neuronal storage disease[57]. To date, megaloneurite is consistently found in the dog[9], monkey[10] and now verified in human In our study. We should point out a limitation, tissues of our human specimen were no longer suitable for NADPH-d staining. We made the conclusion from examination of VIP immunoreactivity, because we found that megaloneurites colocalize with VIP in both dog and non-human primates.

Distribution of VIP immunopositive fibers and terminals mainly occur in the lumbosacral spinal cord in human[12] and animals[58, 59]. Swelling fibers are usually considered as the neurodegenerative pathology[60-62]. Meanwhile, similar morphological assessment terms as neurite swelling or neuronal dystrophy[63-65]. Part of the neurite swelling shows a relative short length[65-67]. The enlarged diameter fibers (Swelling or dystrophy) could be caused by neurotoxic[63, 68], physical damage[64, 69, 70] and aging[71, 72] as well as genetic disease[64, 65]. One of a specialized neurodegeneration is considered for lysosomal storage disease[67]. It shows axon spheroid and abnormal swellings located proximal to initial segments of soma in cerebral cortex[67]. Megaloneurites was anatomical depended in the sacral spinal cord of aged dog, monkeys and human. Our finding of aging-related NADPH-d body in aged rat occurs in the lumbosacral spinal cord[73], which is a spheroidal formation. The megaloneurites, the aging-related alteration will indicated the dysfunction of pelvic organs, because VIP innervate the human pelvic organs, such as vagina[74], urinary bladder[75] and other the genitourinary system of man and animals[76]. VIP is a neurotransmitter and or constitutive component in afferent projections from the pelvic viscera[77-79]. VIP-ergic fibers innervate urogenital organs[5, 14, 24, 80-82]. In major pelvic ganglia, VIP and NOS are found co-localized in a part of small-sized neurons[83]. Immunohistochemistry of VIP reveals that the localization is primarily most prominent in afferent axons and terminals in Lissauer’s tract and in lateral laminae I and V of the dorsal horn in sacral segments of the cat’s spinal cord[58]. Many VIP neurons exhibit positive co-localization with an androgen receptor nucleus[84]. Interestingly, different pelvic organs may be innervated by a convergent pathways[85]. Coordinate sexual reflexes may also need integrated central regulation pathways in spinal cord and brain[86]. VIP may be relevant to the neural control of micturition[87, 88], constipation[89], erectile function[23, 79, 90]. For study of canine penile erection, VIP may play a potential role in canine penile erection[20-22, 79]. NO-ergic neurons are identified in the spinal cord[91]. As mentioned above, NADPH-d staining is associated to identify the NOS neuron[28, 91]. NOS-ergic neurons are distributed in the spinal cord[92]. NOS-ergic and NADPH-d-ergic neurons also innervate the pelvic organs[93, 94]. Similarly, NOS and NADPH-d neurons also innervate pelvic organs and play important role for regulation of urogenital functions[95-98]. Aging-related alteration may cause impaired function of pelvic organs.

We thought that VIP megaloneurites were aging-related neurodegenerative neuropathology and specialized in the sacral spinal cord[9, 10]. Thick VIP fascicle and many thin varicose VIP fibers are detected in the intermediate gray[8]. But it is not theoretically and systemically investigated the distribution and discussed aging-related characteristic and other neurochemical property. Almost all of megaloneurites distributed not only occurred in the grey matter and also in the white matter. Beside NADPH-d and NOS megaloneurite, it is important for the occurrence of VIP megaloneurite in lateral funiculus of white matter in aged human sacral spinal cord. Paraventricular hypothalamic nucleus sends descending projection through the lateral funiculus to sacral spinal cord[99]. It corelates to the intermediolateral cell column of the thoracic spinal cord and the sacral parasympathetic nucleus and DGC[99]. In human, functional connection to up cervical spinal segments can be tested in the region of dorsal-lateral funiculus of lumbosacral spinal cord[100]. Some of the sacral spinal neurons send projection through the lateral funiculus to contralateral side to up level of spinal cord[101]. We presumed that megaloneurites could cause dysfunction of visceral motoneurons. The distribution of megaloneurites is limited within the sacral spinal cord[9, 10]. Propriospinal morphological function was supposedly tested by lesion of local lateral funiculus[102, 103]. But we did not find VIP positive cell bodies in the sacral spinal cord, the VIP megaloneurite axonal course located within lateral funiculus. The results suggested that VIP positive somas located outside of spinal cord. According to our previous study, the course of megaloneurites oriented two directions, coronal and horizontal orientations[9, 10]. Definitely, megaloneurites distributed two different compartments in the sacral spinal cord: gray matter and white matter. The numbers of axons in lateral funiculus in rat has been study in the sacral spinal cord[104]. We should consider the measurement of aging study in future.

Deposition of α-synuclein in the LCP, a region of the sacral spinal dorsal horn important for the relay of pelvic visceral afferents. We also did examination of thoracic level of human spinal cord(data not showed here). We noted no significant change of the expression of α-synuclein in spinal cord between young and aged dog[105]. In this study, they do not test the sacral spinal cord by using western blot analysis of α-synulein protein level between the tissues of adult and aged dogs. Actually, accumulation of α-synuclein causes substantial changes[106]. Our experiment focused on the morphological examination in the sacral spinal cord of aged human. In our present study, examination of α-synuclein immunoreactivity provided a background control to some extent. The aging of sacral spinal cord should be considered as a risk factor for bladder and bowel dysfunction[107].The number of the patients for medical treatment will predict increase in future[108, 109]. Aging animal model should be more important for urogenital dysfunction. In general, dog is suggested to be suitable animal model for aging research [36, 37]. These previous dog model of urogenital dysfunction should be considered with aging study in future[110, 111]. In present study, biochemical and immunoreactive property of megaloneurites were relevant to NAHPH-d, VIP and NOS occurred in the sacral spinal cord of aged dog. At least, VIP megaloneurites was evidenced in the sacral spinal cord of aged human. The identification of megaloneurites provided a basis for a later study on the aged sacral spinal cord. Further experimental study, neuronal tracing and two-dimensional gel electrophoresis combined with protein identification by mass spectrometry should be considered for proteomics.

VIP megaloneurites in aged non-human primates is consistently with that of aged dog[9, 10] and verified in the sacral spinal cord of aged human. The sacral spinal cord of aged dog should be suggested for aging urogenital disorders of human being. Considered with distribution of NADPH-d histology, NOS and VIP immunoreactivity in the pelvic organs and the localization in spinal cord, aging-related megaloneurites implicate the aging-related dysfunction of pelvic organs.

## Acknowledgments

This work was supported by grants from National Natural Science Foundation of China (81471286), Undergraduate Training Programs for Innovation and Entrepreneurship of Liaoning (201410160007) and Research Start-Up Grant for New Science Faculty of Jinzhou Medical University (173514017).

## Author Contributions

YL,HT,WH,YJ and ZW conceived and performed the experiments as well as analyzed data. YJ, WH, CR, ZW, XX, HL, FL, XW, TZ, HJ, JS, HT assisted YL and ZW in management of lab. HT provided important experimental guidance. YL and HT prepared the figures. HT, YL, WH, XW, TZ, CR and ZW discussed the results. HT wrote the manuscript. HT supervised the project and coordinated the study.

## References

[1] G. Deng, L. Jin, The effects of vasoactive intestinal peptide in neurodegenerative disorders, Neurol Res, 39 (2017) 65–72.

[2] D.E. Brenneman, T.M. Phillips, B.W. Festoff, I. Gozes, Identity of neurotrophic molecules released from astroglia by vasoactive intestinal peptide, Ann N Y Acad Sci, 814 (1997) 167–173.

[3] Y. Iwasaki, K. Ikeda, Y. Ichikawa, O. Igarashi, Vasoactive intestinal peptide influences neurite outgrowth in cultured rat spinal cord neurons, Neurol Res, 23 (2001) 851–854.

[4] A.I. Basbaum, E.J. Glazer, Immunoreactive vasoactive intestinal polypeptide is concentrated in the sacral spinal cord: a possible marker for pelvic visceral afferent fibers, Somatosensory research, 1 (1983) 69–82.

[5] M. Kawatani, I.P. Lowe, I. Nadelhaft, C. Morgan, W.C. De Groat, Vasoactive intestinal polypeptide in visceral afferent pathways to the sacral spinal cord of the cat, Neuroscience letters, 42 (1983) 311–316.

[6] J.M. Lundberg, T. Hökfelt, Ö. Nilsson, L. Terenius, J. Rehfeld, R. Elde, S. Sald, Peptide neurons in the vagus, splanchnic and sciatic nerves*, Acta Physiologica Scandinavica, 104 (1978) 499–501.

[7] J.R. Keast, W.C. De Groat, Segmental distribution and peptide content of primary afferent neurons innervating the urogenital organs and colon of male rats, The Journal of comparative neurology, 319 (1992) 615–623.

[8] K. Chung, R.P. Briner, S.M. Carlton, K.N. Westlund, Immunohistochemical localization of seven different peptides in the human spinal cord, The Journal of comparative neurology, 280 (1989) 158–170.

[9] Y. Li, Y. Jia, W. Hou, Z. Wei, X. Wen, Y. Tian, W. Zhang, L. Bai, A. Guo, G. Du, H. Tan, De novo aging-related megaloneurites: alteration of NADPH diaphorase positivity in the sacral spinal cord of the aged dog, bioRxiv, DOI 10.1101/483990(2019) 483990.

[10] Y. Li, Z. Wei, Y. Jia, W. Hou, Y. Wang, S. Yu, G. Shi, G. Du, H. Tan, Dual forms of aging-related NADPH diaphorase neurodegeneration in the sacral spinal cord of aged non-human primates, bioRxiv, DOI 10.1101/527358(2019)527358.

[11] E.E. Benarroch, L.D. Aimone, T.L. Yaksh, Segmental analysis of neuropeptide concentrations in normal human spinal cord, Neurology, 40 (1990) 137–144.

[12] P. Anand, S.J. Gibson, G.P. McGregor, M.A. Blank, M.A. Ghatei, A.J. Bacarese-Hamilton, J.M. Polak, S.R. Bloom, A VIP-containing system concentrated in the lumbosacral region of human spinal cord, Nature, 305 (1983) 143–145.

[13] Y. Charnay, J.A. Chayvulle, S.I. Said, P.M. Dubois, Localization of vasoactive intestinal peptide immunoreactivity in human foetus and newborn infant spinal cord, Neuroscience, 14 (1985) 195–205.

[14] J.D. Tompkins, B.M. Girard, M.A. Vizzard, R.L. Parsons, VIP and PACAP effects on mouse major pelvic ganglia neurons, Journal of molecular neuroscience: MN, 42 (2010) 390–396.

[15] R. Crowe, K. Light, C.P. Chilton, G. Burnstock, Vasoactive intestinal polypeptide-, somatostatin- and substance P-immunoreactive nerves in the smooth and striated muscle of the intrinsic external urethral sphincter of patients with spinal cord injury, The Journal of urology, 136 (1986) 487–491.

[16] J. Gu, J.M. Polak, L. Probert, K.N. Islam, P.J. Marangos, S. Mina, T.E. Adrian, G.P. Mcgregor, D.J.O. Shaughnessy, S.R. Bloom, Peptidergic Innervation of the Human Male Genital Tract, The Journal of urology, 130 (1983) 386–391.

[17] J.M. Polak, S.R. Bloom, Localisation and measurement of VIP in the genitourinary system of man and animals, Peptides, 5 (1984) 225–230.

[18] G. Ishiyama, N. Hinata, Y. Kinugasa, G. Murakami, M. Fujimiya, Nerves supplying the internal anal sphincter: an immunohistochemical study using donated elderly cadavers, Surgical and Radiologic Anatomy, 36 (2014) 1033–1042.

[19] P. Biancani, J. Walsh, J. Behar, Vasoactive intestinal peptide: A neurotransmitter for relaxation of the rabbit internal anal sphincter*, Gastroenterology, 89 (1985) 867–874.

[20] P.O. Andersson, S.R. Bloom, S. Mellander, Haemodynamics of pelvic nerve induced penile erection in the dog: possible mediation by vasoactive intestinal polypeptide, The Journal of Physiology, 350 (1984) 209–224.

[21] H. Aoki, J. Matsuzaka, K. Yeh, F. Sato, T. Fujioka, T. Kubo, T. Ohhori, N. Yasuda, Involvement of Vasoactive Intestinal Peptide (VIP) as a Humoral Mediator of Penile Erectile Function in the Dog, Journal of Andrology, 15 (1994) 174–182.

[22] Y. Takahashi, S.R. Aboseif, F. Benard, C.G. Stief, T.F. Lue, E.A. Tanagho, Effect of Intracavernous Simultaneous Injection of Acetylcholine and Vasoactive Intestinal Polypeptide on Canine Penile Erection, The Journal of urology, 148 (1992) 446–448.

[23] J. Gu, M. Lazarides, J.P. Pryor, M.A. Blank, J.M. Polak, R. Morgan, P.J. Marangos, S.R. Bloom, DECREASE OF VASOACTIVE INTESTINAL POLYPEPTIDE (VIP) IN THE PENISES FROM IMPOTENT MEN, The Lancet, 324 (1984) 315–318.

[24] L. Zhu, J. Lang, F. Jiang, X. Jiang, J. Chen, Vasoactive intestinal peptide in vaginal epithelium of patients with pelvic organ prolapse and stress urinary incontinence, International Journal of Gynecology & Obstetrics, 105 (2009) 223–225.

[25] H.E. Bredkjoer, C. Palle, E. Ekblad, J. Fahrenkrug, B. Ottesen, PreproVIP-derived peptides in the human female genital tract: expression and biological function, Neuropeptides, 31 (1997) 209–215.

[26] A. Gallo, M. Leerink, B. Michot, E. Ahmed, P. Forget, A. Mouraux, E. Hermans, R. Deumens, Bilateral tactile hypersensitivity and neuroimmune responses after spared nerve injury in mice lacking vasoactive intestinal peptide, Exp Neurol, 293 (2017) 62–73.

[27] P. Anand, M.A. Ghatei, N.D. Christofides, M.A. Blank, G.P. McGregor, J.F. Morrison, F. Scaravilli, S.R. Bloom, Differential neuropeptide expression after visceral and somatic nerve injury in the cat and rat, Neuroscience letters, 128 (1991) 57–60.

[28] B.T. Hope, G.J. Michael, K.M. Knigge, S.R. Vincent, Neuronal NADPH diaphorase is a nitric oxide synthase, Proceedings of the National Academy of Sciences of the United States of America, 88 (1991) 2811–2814.

[29] K.Y. Yeh, C.H. Wu, Y.F. Tsai, Noncontact erection is enhanced by Ginkgo biloba treatment in rats: role of neuronal NOS in the paraventricular nucleus and sacral spinal cord, Psychopharmacology (Berl), 222 (2012) 439–446.

[30] A. Kisucka, L. Hricova, J. Pavel, J.B. Strosznajder, M. Chalimoniuk, J. Langfort, J. Galik, M. Marsala, J. Radonak, N. Lukacova, Baclofen or nNOS inhibitor affect molecular and behavioral alterations evoked by traumatic spinal cord injury in rat spinal cord, The spine journal: official journal of the North American Spine Society, 15 (2015) 1366–1378.

[31] S. Gao, C. Cheng, J. Zhao, M. Chen, X. Li, S. Shi, S. Niu, J. Qin, M. Lu, A. Shen, Developmental regulation of PSD-95 and nNOS expression in lumbar spinal cord of rats, Neurochemistry international, 52 (2008) 495–501.

[32] M.A. Vizzard, K. Erickson, W.C. de Groat, Localization of NADPH diaphorase in the thoracolumbar and sacrococcygeal spinal cord of the dog, Journal of the autonomic nervous system, 64 (1997) 128–142.

[33] N. Lukacova, J. Kafka, D. Cizkova, M. Marsala, J. Marsala, The effect of cauda equina constriction on nitric oxide synthase activity, Neurochemical research, 29 (2004) 429–439.

[34] A.H. Pullen, P. Humphreys, R.G. Baxter, Comparative analysis of nitric oxide synthase immunoreactivity in the sacral spinal cord of the cat, macaque and human, Journal of anatomy, 191 (Pt 2) (1997) 161–175.

[35] M.G. Ferrini, T.R. Magee, D. Vernet, J. Rajfer, N.F. Gonzalez-Cadavid, Penile neuronal nitric oxide synthase and its regulatory proteins are present in hypothalamic and spinal cord regions involved in the control of penile erection, The Journal of comparative neurology, 458 (2003) 46–61.

[36] K.M. Gilmore, K.A. Greer, Why is the dog an ideal model for aging research?, Experimental gerontology, 71 (2015) 14–20.

[37] J.M. Hoffman, K.E. Creevy, A. Franks, D.G. O’Neill, D.E.L. Promislow, The companion dog as a model for human aging and mortality, Aging Cell, 17 (2018) e12737.

[38] J. Orendacova, M. Marsala, I. Sulla, J. Kafka, P. Jalc, D. Cizkova, Y. Taira, J. Marsala, Incipient cauda equina syndrome as a model of somatovisceral pain in dogs: spinal cord structures involved as revealed by the expression of c-fos and NADPH diaphorase activity, Neuroscience, 95 (2000) 543–557.

[39] M. Oinas, A. Paetau, L. Myllykangas, I.L. Notkola, H. Kalimo, T. Polvikoski, alpha-Synuclein pathology in the spinal cord autonomic nuclei associates with alpha-synuclein pathology in the brain: a population-based Vantaa 85+ study, Acta neuropathologica, 119 (2010) 715–722.

[40] V.G. VanderHorst, T. Samardzic, C.B. Saper, M.P. Anderson, S. Nag, J.A. Schneider, D.A. Bennett, A.S. Buchman, α-synuclein pathology accumulates in sacral spinal visceral sensory pathways, Annals of neurology, 78 (2015) 142–149.

[41] H. Sumikura, M. Takao, H. Hatsuta, S. Ito, Y. Nakano, A. Uchino, A. Nogami, Y. Saito, H. Mochizuki, S. Murayama, Distribution of alpha-synuclein in the spinal cord and dorsal root ganglia in an autopsy cohort of elderly persons, Acta neuropathologica communications, 3 (2015) 57.

[42] C. Morgan, I. Nadelhaft, W.C. de Groat, The distribution of visceral primary afferents from the pelvic nerve to Lissauer’s tract and the spinal gray matter and its relationship to the sacral parasympathetic nucleus, The Journal of comparative neurology, 201 (1981) 415–440.

[43] B.F.M. Blok, J.T.P.W. Van Maarseveen, G. Holstege, Electrical stimulation of the sacral dorsal gray commissure evokes relaxation of the external urethral sphincter in the cat, Neuroscience letters, 249 (1998) 68–70.

[44] C.A. Sasek, V.S. Seybold, R. Elde, The immunohistochemical localization of nine peptides in the sacral parasympathetic nucleus and the dorsal gray commissure in rat spinal cord, Neuroscience, 12 (1984) 855–873.

[45] A.I. Basbaum, E.J. Glazer, Immunoreactive Vasoactive Intestinal Polypeptide Is Concentrated in the Sacral Spinal Cord: A Possible Marker for Pelvic Visceral Afferent Fibers, Somatosensory and Motor Research, 1 (1983) 69–82.

[46] W.C. De Groat, Neuropeptides in pelvic afferent pathways, Cellular and Molecular Life Sciences, 43 (1987) 801–813.

[47] X. Zhang, G. Ju, R. Elde, T. Hokfelt, Effect of peripheral nerve cut on neuropeptides in dorsal root ganglia and the spinal cord of monkey with special reference to galanin, J Neurocytol, 22 (1993) 342–381.

[48] J.M. Allen, S.J. Gibson, T.E. Adrian, J.M. Polak, S.R. Bloom, Neuropeptide Y in human spinal cord, Brain Research, 308 (1984) 145–148.

[49] T. Hokfelt, X. Zhang, Z. Wiesenfeldhallin, Messenger plasticity in primary sensory neurons following axotomy and its functional implications, Trends in Neurosciences, 17 (1994) 22–30.

[50] L.L. Scharer, H.J. Burhenne, Megacolon, a nonspecific sign: clinical classification and roentgenologic differentiation, Radiologia clinica, 34 (1965) 236.

[51] M.F. Campbell, Primary Megalo-Ureter, The Journal of urology, 68 (1952) 584–590.

[52] K.P. Dikshit, G.P. Mathur, I.P. Elhence, Congenital megalo-urethra with giantism of the penis, Indian Journal of Pediatrics, 42 (1975) 27–28.

[53] G. Joe, M. Andoh, A. Shinoki, W. Lang, Y. Kumagai, J. Sadahiro, M. Okuyama, A. Kimura, H. Shimizu, H. Hara, Megalo-type α-1,6-glucosaccharides induce production of tumor necrosis factor α in primary macrophages via toll-like receptor 4 signaling, Biomedical Research-tokyo, 37 (2016) 179–186.

[54] B. Mulloney, Structure of the Giant Fibers of Earthworms, Science, 168 (1970) 994–996.

[55] M.J. Allen, T.A. Godenschwege, Electrophysiological Recordings from the Drosophila Giant Fiber System (GFS), CSH Protocols, 2010 (2010).

[56] A.P. Yong, E. Bednarski, C.M. Gall, G. Lynch, C.E. Ribak, Lysosomal dysfunction results in lamina-specific meganeurite formation but not apoptosis in frontal cortex, Exp Neurol, 157 (1999) 150–160.

[57] S.U. Walkley, L.F. James, Locoweed-induced neuronal storage disease characterized by meganeurite formation, Brain Res, 324 (1984) 145–150.

[58] M. Kawatani, I.P. Lowe, I. Nadelhaft, C. Morgan, W.C. De Groat, Vasoactive intestinal polypeptide in visceral afferent pathways to the sacral spinal cord of the cat, Neuroscience letters, 42 (1983) 311–316.

[59] W.C. de Groat, M. Kawatani, T. Hisamitsu, I. Lowe, C. Morgan, J. Roppolo, A.M. Booth, I. Nadelhaft, D. Kuo, K. Thor, The role of neuropeptides in the sacral autonomic reflex pathways of the cat, Journal of the autonomic nervous system, 7 (1983) 339–350.

[60] B. Veronesi, E.R. Peterson, M.B. Bornstein, P.S. Spencer, Ultrastructural studies of the dying-back process. VI. Examination of nerve fibers undergoing giant axonal degeneration in organotypic culture, Journal of neuropathology and experimental neurology, 42 (1983) 153–165.

[61] M. Schuelke, J. Cervos-Navarro, Degenerative changes in unmyelinated nerve fibers in late-infantile neuronal ceroidlipofuscinosis. A morphometric study of conjunctival biopsy specimens, Acta neuropathologica, 95 (1998) 175–183.

[62] P.S. Spencer, H.H. Schaumburg, Ultrastructural studies of the dying-back process. IV. Differential vulnerability of PNS and CNS fibers in experimental central-peripheral distal axonopathies, Journal of neuropathology and experimental neurology, 36 (1977) 300–320.

[63] C.E. Menard, M. Durston, E. Zherebitskaya, D.R. Smith, D. Freed, G.W. Glazner, G. Tian, P. Fernyhough, R.C. Arora, Temporal dystrophic remodeling within the intrinsic cardiac nervous system of the streptozotocin-induced diabetic rat model, Acta neuropathologica communications, 2 (2014) 60.

[64] K.L. Newell, P. Boyer, E. Gomez-Tortosa, W. Hobbs, E.T. Hedley-Whyte, J.P. Vonsattel, B.T. Hyman, Alpha-synuclein immunoreactivity is present in axonal swellings in neuroaxonal dystrophy and acute traumatic brain injury, Journal of neuropathology and experimental neurology, 58 (1999) 1263–1268.

[65] A. Tarrade, C. Fassier, S. Courageot, D. Charvin, J. Vitte, L. Peris, A. Thorel, E. Mouisel, N. Fonknechten, N. Roblot, D. Seilhean, A. Dierich, J.J. Hauw, J. Melki, A mutation of spastin is responsible for swellings and impairment of transport in a region of axon characterized by changes in microtubule composition, Human molecular genetics, 15 (2006) 3544–3558.

[66] S.U. Walkley, S. Wurzelmann, D.P. Purpura, Ultrastructure of neurites and meganeurites of cortical pyramidal neurons in feline gangliosidosis as revealed by the combined Golgi-EM technique, Brain Res, 211 (1981) 393–398.

[67] S.U. Walkley, A.L. Pierok, Ferric ion-ferrocyanide staining in ganglioside storage disease establishes that meganeurites are of axon hillock origin and distinct from axonal spheroids, Brain Res, 382 (1986) 379–386.

[68] A. Simonati, N. Rizzuto, J.B. Cavanagh, The effects of 2,5-hexanedione on axonal regeneration after nerve crush in the rat, Acta neuropathologica, 59 (1983) 216–224.

[69] D.G. Emery, J.H. Lucas, G.W. Gross, The sequence of ultrastructural changes in cultured neurons after dendrite transection, Experimental brain research, 67 (1987) 41–51.

[70] F.T. Sayer, M. Oudega, T. Hagg, Neurotrophins reduce degeneration of injured ascending sensory and corticospinal motor axons in adult rat spinal cord, Exp Neurol, 175 (2002) 282–296.

[71] O. Bugiani, G. Giaccone, L. Verga, B. Pollo, B. Ghetti, B. Frangione, F. Tagliavini, Alzheimer patients and Down patients: abnormal presynaptic terminals are related to cerebral preamyloid deposits, Neuroscience letters, 119 (1990) 56–59.

[72] R.E. Schmidt, L. Beaudet, S.B. Plurad, W.D. Snider, K.G. Ruit, Pathologic alterations in pre- and postsynaptic elements in aged mouse sympathetic ganglia, J Neurocytol, 24 (1995) 189–206.

[73] H. Tan, J. He, S. Wang, K. Hirata, Z. Yang, A. Kuraoka, M. Kawabuchi, Age-related NADPH-diaphorase positive bodies in the lumbosacral spinal cord of aged rats, Archives of histology and cytology, 69 (2006) 297–310.

[74] C.H. Hoyle, R.W. Stones, T. Robson, K. Whitley, G. Burnstock, Innervation of vasculature and microvasculature of the human vagina by NOS and neuropeptide-containing nerves, Journal of anatomy, 188 (Pt 3) (1996) 633–644.

[75] R. Crowe, J. Vale, K.R. Trott, P. Soediono, T. Robson, G. Burnstock, Radiation-induced changes in neuropeptides in the rat urinary bladder, The Journal of urology, 156 (1996) 2062–2066.

[76] J.M. Polak, S.R. Bloom, Localisation and measurement of VIP in the genitourinary system of man and animals, Peptides, 5 (1984) 225–230.

[77] K. Juenemann, T.F. Lue, J. Luo, S.A. Jadallah, L. Nunes, E.A. Tanagho, The Role of Vasoactive Intestinal Polypeptide as a Neurotransmitter in Canine Penile Erection: A Combined in Vivo and Immunohistochemical Study, The Journal of urology, 138 (1987) 871–877.

[78] K.-P. Jünemann, T.F. Lue, H. Melchior, E.A. Tanagho, VIP — A Peripheral Neurotransmitter in Penile Erection, Springer Berlin Heidelberg, Berlin, Heidelberg, 1987, pp. 163–166.

[79] B. Ottesen, G. Wagner, R. Virag, J. Fahrenkrug, Penile erection: possible role for vasoactive intestinal polypeptide as a neurotransmitter, BMJ, 288 (1984) 9–11.

[80] M. Botti, L. Ragionieri, A. Cacchioli, R. Panu, F. Gazza, Immunohistochemical Properties of the Peripheral Neurons Projecting to the Pig Bulbospongiosus Muscle, Anatomical record (Hoboken, N.J.: 2007), 299 (2016) 1192–1202.

[81] Z. Grozdanovic, H.G. Baumgarten, Colocalisation of NADPH-diaphorase with neuropeptides in the ureterovesical ganglia of humans, Acta Histochem, 98 (1996) 245–253.

[82] J. Gu, J.M. Polak, H.C. Su, M.A. Blank, J.F. Morrison, S.R. Bloom, Demonstration of paracervical ganglion origin for the vasoactive intestinal peptide-containing nerves of the rat uterus using retrograde tracing techniques combined with immunocytochemistry and denervation procedures, Neuroscience letters, 51 (1984) 377–382.

[83] T. Domoto, T. Tsumori, Co-localization of nitric oxide synthase and vasoactive intestinal peptide immunoreactivity in neurons of the major pelvic ganglion projecting to the rat rectum and penis, Cell Tissue Res, 278 (1994) 273–278.

[84] J.R. Keast, R.J. Saunders, Testosterone has potent, selective effects on the morphology of pelvic autonomic neurons which control the bladder, lower bowel and internal reproductive organs of the male rat, Neuroscience, 85 (1998) 543–556.

[85] J.A. Christianson, R. Liang, E.E. Ustinova, B.M. Davis, M.O. Fraser, M.A. Pezzone, Convergence of bladder and colon sensory innervation occurs at the primary afferent level, Pain, 128 (2007) 235–243.

[86] A.D. Dobberfuhl, T. Oti, H. Sakamoto, L. Marson, Identification of CNS Neurons Innervating the Levator Ani and Ventral Bulbospongiosus Muscles in Male Rats, The Journal of Sexual Medicine, 11 (2014) 664–677.

[87] S. Studeny, B.P. Cheppudira, S. Meyers, E.M. Balestreire, G. Apodaca, L.A. Birder, K.M. Braas, J.A. Waschek, V. May, M.A. Vizzard, Urinary Bladder Function and Somatic Sensitivity in Vasoactive Intestinal Polypeptide (VIP)-/-Mice, Journal of Molecular Neuroscience, 36 (2008) 175–187.

[88] J. Gu, J.M. Restorick, M.A. Blank, W.M. Huang, J.M. Polak, S.R. Bloom, A.R. Mundy, Vasoactive intestinal polypeptide in the normal and unstable bladder, BJUI, 55 (1983) 645–647.

[89] T.R. Koch, J.A. Carney, L. Go, V.L.W. Go, Idiopathic chronic constipation is associated with decreased colonic vasoactive intestinal peptide, Gastroenterology, 94 (1988) 300–310.

[90] M. Zhang, Z. Shen, C. Zhang, W. Wu, P. Gao, S. Chen, W. Zhou, Vasoactive intestinal polypeptide, an erectile neurotransmitter, improves erectile function more significantly in castrated rats than in normal rats, BJUI, 108 (2011) 440–446.

[91] J.G. Valtschanoff, R.J. Weinberg, A. Rustioni, NADPH diaphorase in the spinal cord of rats, The Journal of comparative neurology, 321 (1992) 209–222.

[92] N.J. Dun, S.L. Dun, U. Forstermann, L.F. Tseng, Nitric oxide synthase immunoreactivity in rat spinal cord, Neuroscience letters, 147 (1992) 217–220.

[93] S. Carrier, P. Zvara, L. Nunes, N.W. Kour, J. Rehman, T.F. Lue, Regeneration of Nitric Oxide Synthase-Containing Nerves After Cavernous Nerve Neurotomy in the Rat, The Journal of urology, 153 (1995) 1722–1727.

[94] J.R. Keast, A possible neural source of nitric oxide in the rat penis, Neuroscience letters, 143 (1992) 69–73.

[95] A. Schirar, F. Giuliano, O. Rampin, J.P. Rousseau, A large proportion of pelvic neurons innervating the corpora cavernosa of the rat penis exhibit NADPH-diaphorase activity, Cell Tissue Res, 278 (1994) 517–525.

[96] A.L. Burnett, S. Saito, M.P. Maguire, H. Yamaguchi, T.S. Chang, D.F. Hanley, Localization of nitric oxide synthase in spinal nuclei innervating pelvic ganglia, The Journal of urology, 153 (1995) 212–217.

[97] H. Yan, J.R. Keast, Neurturin regulates postnatal differentiation of parasympathetic pelvic ganglion neurons, initial axonal projections, and maintenance of terminal fields in male urogenital organs, The Journal of comparative neurology, 507 (2008) 1169–1183.

[98] J. Rajfer, W.J. Aronson, P.A. Bush, F.J. Dorey, L.J. Ignarro, Nitric Oxide as a Mediator of Relaxation of the Corpus Cavernosum in Response to Nonadrenergic, Noncholinergic Neurotransmission, The New England journal of medicine, 326 (1992) 90–94.

[99] Z. Wiesenfeldhallin, T. Hokfelt, J.M. Lundberg, W.G. Forssmann, M. Reinecke, F.A. Tschopp, J.A. Fischer, Immunoreactive calcitonin gene-related peptide and substance P coexist in sensory neurons to the spinal cord and interact in spinal behavioral responses of the rat, Neuroscience letters, 52 (1984) 199–204.

[100] D. Jeanmonod, M. Sindou, F. Mauguiere, The human cervical and lumbo-sacral evoked electrospinogram. Data from intra-operative spinal cord surface recordings, Electroencephalography and Clinical Neurophysiology, 80 (1991) 477–489.

[101] K. Grottel, D. Bukowska, J. Huber, J. Celichowski, Distribution of the sacral neurones of origin of the ascending spinal tracts with axons passing through the lateral funiculi of the lowermost thoracic segments: an experimental HRP study in the cat, Neuroscience Research, 34 (1999) 67–72.

[102] A. Rustioni, H.G.J.M. Kuypers, G. Holstege, Propriospinal projections from the ventral and lateral funiculi to the motoneurons in the lumbosacral cord of the cat, Brain Research, 34 (1971) 255–275.

[103] I. Molenaar, A. Rustioni, H.G.J.M. Kuypers, The location of cells of origin of the fibers in the ventral and the lateral funiculus of the cat’s lumbo-sacral cord, Brain Research, 78 (1974) 239–254.

[104] K. Chung, R.E. Coggeshall, Numbers of axons in lateral and ventral funiculi of rat sacral spinal cord, The Journal of comparative neurology, 214 (1983) 72–78.

[105] J.H. Ahn, J.H. Choi, J.H. Park, B.C. Yan, I.H. Kim, J.C. Lee, D.H. Lee, J.S. Kim, H.C. Shin, M.H. Won, Comparison of alpha-synuclein immunoreactivity in the spinal cord between the adult and aged beagle dog, Lab Anim Res, 28 (2012) 165–170.

[106] S. Mendritzki, S. Schmidt, T. Sczepan, X.R. Zhu, D. Segelcke, H. Lubbert, Spinal cord pathology in alpha-synuclein transgenic mice, Parkinson’s disease, 2010 (2010) 375462.

[107] R.N. Ranson, M.J. Saffrey, Neurogenic mechanisms in bladder and bowel ageing, Biogerontology, 16 (2015) 265–284.

[108] J.M. Wu, A. Kawasaki, A.F. Hundley, A.A. Dieter, E.R. Myers, V.W. Sung, Predicting the number of women who will undergo incontinence and prolapse surgery, 2010 to 2050, American Journal of Obstetrics and Gynecology, 205 (2011) 230.

[109] J.M. Wu, A.F. Hundley, R.G. Fulton, E.R. Myers, Forecasting the prevalence of pelvic floor disorders in U.S. Women: 2010 to 2050, Obstetrics & Gynecology, 114 (2009) 1278–1283.

[110] D. Newgreen, C. Anderson, P. Nunn, A.J. Carter, M. Bushfield, Models of Urogenital Dysfunction: Urinary Urge Incontinence (UUI) in Anesthetized Dog and Rat Models, Current protocols in pharmacology, 2 (2001).

[111] A.L. Beck, J. Grierson, D.M. Ogden, M.H. Hamilton, V.J. Lipscomb, Outcome of and complications associated with tube cystostomy in dogs and cats: 76 cases (1995-2006), Javma-journal of The American Veterinary Medical Association, 230 (2007) 1184–1189.

